# *In silico* comparative RNA-seq analysis reveals varietal-specific intergenic small open reading frames in *Cucumis sativus* L.

**DOI:** 10.1101/2022.10.13.512045

**Authors:** Gabrielle Shiao Wei Chieng, Boon Chin Tan, Chee How Teo

## Abstract

Small open reading frames (sORFs) have been reported to play important roles in growth, regulation of morphogenesis, and abiotic stress responses in various plant species. However, their sequences and functions remain poorly understood in many plant species including *Cucumis sativus. Cucumis sativus* (commonly known as cucumber) is Asia’s fourth most important vegetable and the second most important crop in Western Europe. The breeding of climate-resilient cucumbers is of great importance to ensure their sustainability under extreme climate conditions. In this study, we aim to isolate the intergenic sORFs from *C. sativus* var. *hardwickii* genome and determine their sequence diversity and expression profiles in *C. sativus* var. *hardwickii* and different cultivars of *C. sativus* var. *sativus* using bioinformatics tools. We identified a total of 50,191 coding sORFs with coding potential (coding sORFs) from *C. sativus* var. *hardwickii* genome. In addition, 1,311 transcribed sORFs were detected in RNA-seq datasets of *C. sativus* var. *hardwickii* and shared homology to sequences deposited in the cucumber EST database, and among these, 91 transcribed sORFs with translation potential were detected. A total of 629 high-confident *C. sativus*-specific sORFs were identified in both varieties. Varietal-specific transcribed sORFs were also identified in *C. sativus* var. *hardwickii* (87) and *C. sativus* var. *sativus* (2,906). Furthermore, cultivar- and tissue-specific transcribed sORFs were identified in different cultivars and tissue samples. The findings of this study provide insight into sequence diversity and expression patterns of sORFs in *C. sativus*, which could help in developing climate-resilient cucumbers.

## 1. Introduction

Cucurbitaceae used to be documented as a monophyletic family without any close relatives (Naegele and Wehner, 2016) but with the addition of more mitochondrial and chloroplast genome sequences of old and new plant materials, a few of the closest relatives were discovered in this family (Chomicki, Schaefer and Renner, 2019). Within Cucurbitaceae, there are roughly 66 species in the genus *Cucumis*, and cucumber (*Cucumis sativus*) is the only one having 2n = 2x = 14 chromosomes (Renner et al., 2007). Cucumber is a valuable crop to humans in terms of its medicinal properties and for the sustenance and economics of certain countries (Mukherjee et al., 2013). Cucumber shows a very high economical value in the fields of therapeutical and cosmetic industries (Mukherjee et al., 2013). Among the many varieties of *Cucumis sativus*, wild cucumber (*C. sativus* var. *hardwickii*), semi-wild Xishuangbanna cucumber (*C. sativus* var. *xishuangbannesis*), the Sikkim cucumber (*C. sativus* var. *sikkimensis*) and the cultivated cucumber (*C. sativus* var. *sativus*) are cross-compatible (Weng, 2021).

The fruits of most wild species are small, bitter, seedy, and non-palatable to humans (Che and Zhang, 2019). After multiple rounds of domestication, Cucurbitaceae crops with higher sugar or carotenoid content, more compact and less branched growth, decreased physical defence, increased fruit size, and non-bitter fruits were created (Chomicki, Schaefer and Renner, 2019). A chemical agent known as cucurbitacin has been reported to exhibit cytotoxicity, anti-inflammatory, anti-fertility agent, and anti-cancer activity (Kaushik et al., 2015). Apart from that, the fruit pulp of cucumber contains a high amount of ascorbic acid, and caffeic acid that helps reduces swelling. As its hard fruit skin is very rich in fibre and several advantageous minerals such as silica, potassium, and magnesium that give a cooling effect, it could be a potential anti-wrinkle agent in the cosmetic industry (Nema et al., 2010).

Today, China has been ranked as one of the top producers and largest domesticators of cucumbers. Based on the statistics from FAOSTAT (2020), the total cucumber production in China in 2018 was 56.24 million tons from 1.044 million hectares. The cucumber production and area utilised for cucumber production in China stand at 52.7% and 74.8% of the corresponding world totals respectively. As for the yield per unit area, China exceeded the world average by 42% with a total of 53.86 kg/ha. Apart from China, India is also a competitive player in terms of its annual cucumber production. In a study by Sanjeev et al. (2017), the annual production of cucumber was 0.698 million tons from 45,000 ha with a productivity of 15.5 t/ha. For both countries, the most concerning issue is having low productivity and challenging climatic diversity (Liu et al., 2020; Sanjeev et al., 2017).

Small open reading frame (sORF) as its name suggests, is a shorter version of the canonical ORF. The size of sORF ranges from 30 bp to 300bp (Hanada et al., 2010). Their minuscule size has caused them to be excluded from most gene prediction methods (Couso and Patraquim, 2017). The length cut-off filter used in most gene prediction methods is 300 bp and any sequences below this cut-off will be considered as being non-functional (Kute et al., 2022). Another contributing factor causing sORF to be diminished is that short sequences normally have evolutionary conservation scores, an indicator of the functionality of a gene (Leong et al., 2022).

To date, there is still no standard classification for sORFs but researchers have made attempts to classify them into different categories (Ong et al., 2022). Ong et al. (2022) have summarized classifications from several studies, namely, sORF (Orr et al., 2020), upstream ORF (uORF; Couso and Patraquim, 2017; Fesenko et al., 2019; Khitun et al., 2019; Orr et al., 2020; Takahashi et al., 2019), intergenic sORF (Couso and Patraquim, 2017; Khitun et al., 2019), sORF with translational potential (Khitun et al., 2019), long noncoding ORF (lncORF; Couso and Patraquim, 2017), short CDS (Couso and Patraquim, 2017), short isoform (Couso and Patraquim, 2017), downstream ORF (dORF; Fesenko et al., 2019), CDS-sORF (Fesenko et al., 2019), interlaced-sORF (Fesenko et al., 2019), miPEP (Takahashi et al., 2019), microprotein (Takahashi et al., 2019), hormone-like peptides (Takahashi et al., 2019), and defensin (Takahashi et al., 2019). Other than coding regions, sORFs can also be found in mitochondrial RNAs, circular RNAs, and long noncoding RNAs (lncRNAs) (Orr et al., 2020).

sORFs have been reported to get translated into small peptides and have functional roles in plants (Castellana et al., 2008; Hanada et al., 2013; Cabrera-Quio et al., 2016). Ong et al. (2022) reviewed the roles of sORF-encoded proteins (SEPs) in several biological processes, including cell signaling, abiotic stress responses, morphogenesis, and growth regulations. In addition, studies also reported that short proteins also function as secreted peptides and hormones. Cabrera-Quio et al. (2016) reported an 18 aa plant polypeptide hormone known as systemin that engages in plant defence mechanisms and secretes phytosulfokine pentapeptides (PSK; 100 aa) that function in regulating plant growth and stress responses.

In this study, we used the reference genome of *C. sativus* var. *hardwickii* PI183967 for *in silico* sORF prediction and characterisation. The main objective of this study was to identify and characterise varietal-specific sORFs from *C. sativus* var. *hardwickii* PI183967 genome. To achieve this objective, we first identify the coding sORFs from the genome sequences of *C. sativus* var. *hardwickii* PI183967 using sORFfinder. We then classified the coding sORFs into coding sORF with transcription potential (transcribed sORF) and with both transcription and translation potential (translated sORF) based on the outcomes from the transcript (RNA-seq and EST) and protein sequence homology search (SWISS-PROT) analysis. We determined the sORF expression profiles in RNA-seq datasets of *C. sativus* var. *hardwickii* and different cultivars of *C. sativus* var. *sativus* using gene expression tools. In addition, we also determined the sORF expression profiles in different tissue samples. Finally, the potential biological functions of the ttsORFs were annotated using Gene Ontology (GO) and the Kyoto Encyclopedia of Genes and Genomes (KEGG) pipeline. The findings from this study will set an important foundation for the development of climate-resilient cucumbers and other crop species.

## 2. Results and Discussion

### Identification of small open reading frame in *C. sativus* var. *hardwickii*

The total number of sORFs in an organism varied from species to species. In *Arabidopsis thaliana*, approximately 33,809 sORFs were predicted from the intergenic regions using sORFfinder with 7,159 sORFs are coding sORFs and 2,996 coding sORFs likely expressed in at least one experimental condition of the tilling array data (Hanada et al., 2007). Using the same sORF detection pipeline, a total of 850,540 sORFs were predicted from the genome of *C. sativus* var. *hardwickii* PI183967 and 50,584 coding sORFs were detected (Table 1). RepeatMasker and CD-HIT were conducted to remove the csORFs with repeat sequence homology and to cluster redundant csORFs to unique coding sORFs. A final set of 50,191 csORFs was blast searched against the Cucumber EST collection to obtain 693 transcribed sORFs that shared high homology to cucumber EST sequences. Out of the 693 transcribed sORFs, 91 showed high homology to protein sequences deposited in the SWISS-PROT database. Using the transcriptome datasets of *C. sativus* var. *hardwickii* and var. *sativus*, we identified 1,311 and 14,799 transcribed sORFs, respectively (Table 1).

**Table 1:**
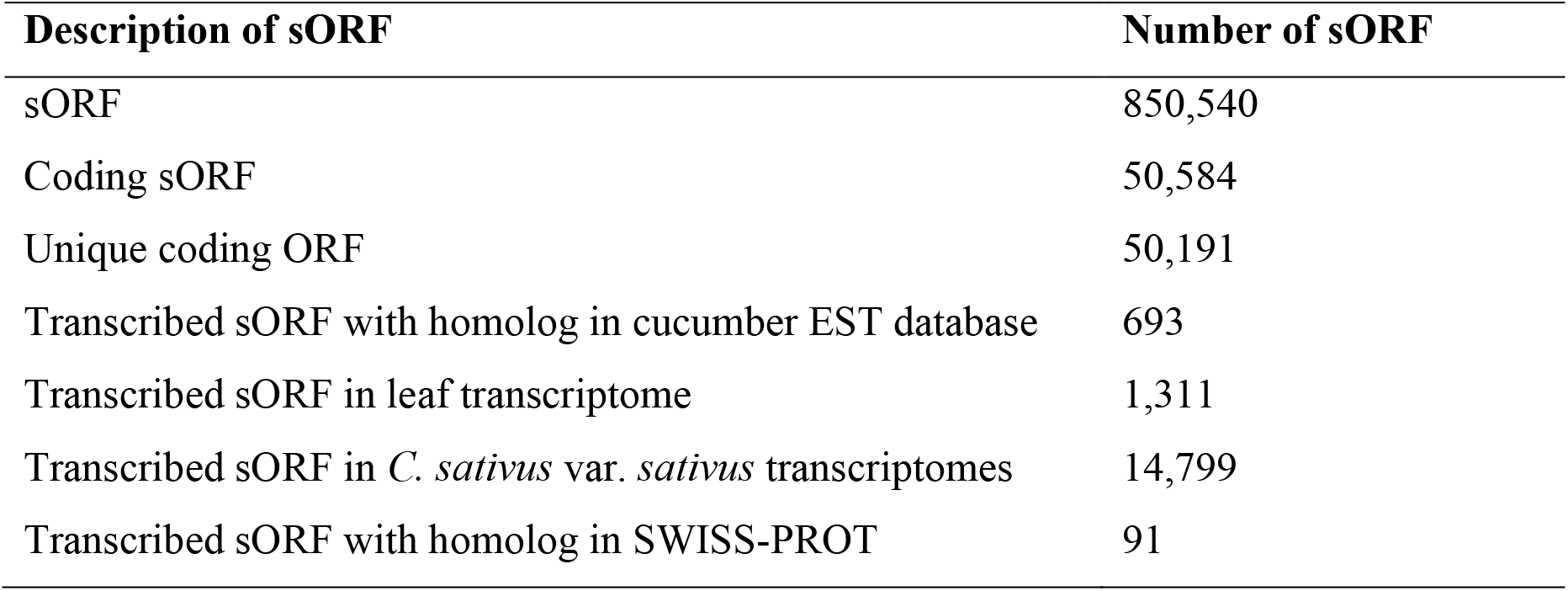
Summary of intergenic small open reading frames found in *C. sativus* var. *hardwickii*.

| Description of sORF | Number of sORF |
| --- | --- |
| sORF | 850,540 |
| Coding sORF | 50,584 |
| Unique coding ORF | 50,191 |
| Transcribed sORF with homolog in cucumber EST database | 693 |
| Transcribed sORF in leaf transcriptome | 1,311 |
| Transcribed sORF in *C. sativus* var. *sativus* transcriptomes | 14,799 |
| Transcribed sORF with homolog in SWISS-PROT | 91 |

### Identification of species- and varietal-specific transcribed sORFs

To find the high confident species-specific transcribed sORFs in *C. sativus* species, a Venn diagram was plotted using the transcribed sORF IDs of *C. sativus* var. *hardwickii* and var. *sativus* (Figure 1). The result showed that there are 629 high confident species-specific transcribed sORFs that can be detected in the transcriptome datasets of *C. sativus* var. *hardwickii* and var. *sativus*. Out of 14,866 transcribed sORFs, 87 were specific to *C. sativus* var. *hardwickii* and 13,555 transcribed sORFs were only found in var. *sativus*. These unique transcribed sORFs were designated as varietal-specific transcribed sORFs. Venn diagram of the transcribed sORFs from 5 different cultivars (Figure 1) also reveals the cultivar-specific transcribed sORFs where cv. Jin You 35 harboured the highest number of cultivars-specific transcribed sORF (2,027), followed by cv. Zaoer-N (1,568), cv. Chinese Long (996) and cv. Jin You (299). Overall, the var. *hardwickii* has the lowest number of transcribed sORF whereas cv. Jin You 35 has the highest (Table 2).

**Figure 1:**
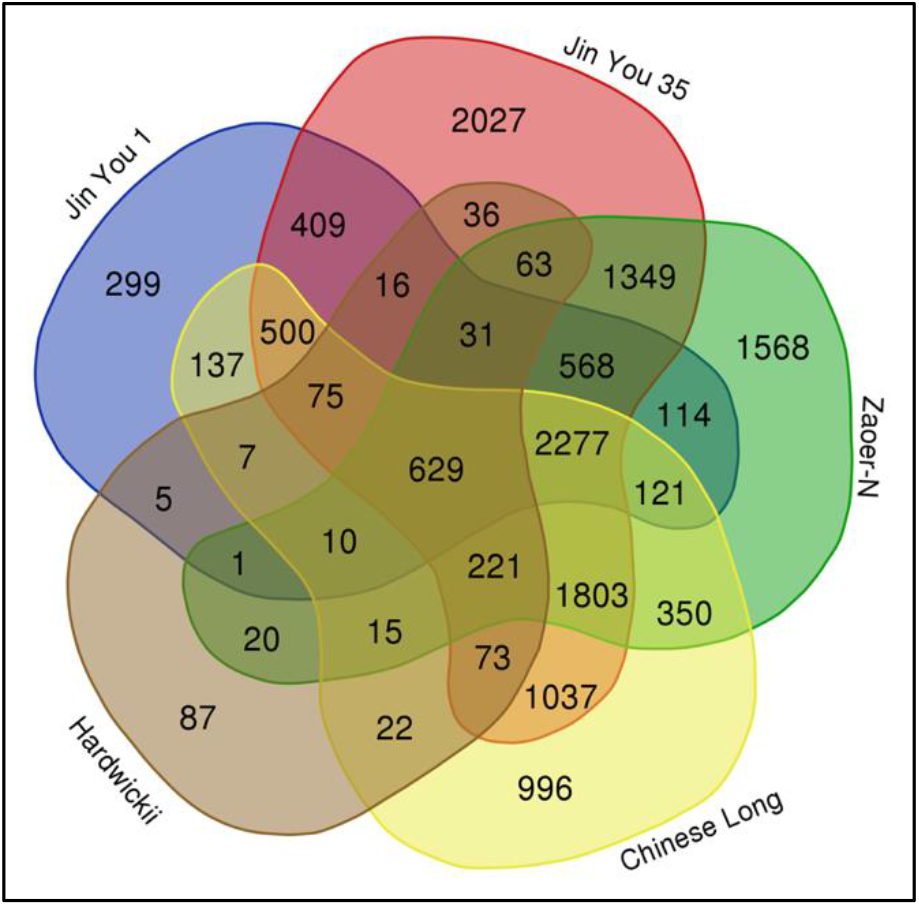
Venn diagram showing the comparison of transcribed sORF in all 5 cultivars.

**Table 2:** Summary of Cucumis sativus var. sativus transcribed sORF and cultivar-specific transcribed sORF.

| Cultivars | Transcribed sORF | Cultivar-specific transcribed sORF |
| --- | --- | --- |
| Jin You 1 | 5,199 | 299 |
| Jin You 35 | 11,114 | 2,027 |
| Zaoer-N | 9,140 | 1,568 |
| Chinese Long | 8,273 | 996 |

### Expression profile of transcribed sORFs in different tissues of *C. sativus*

To study the tissue-specific expression of transcribed sORFs in *C. sativus* var. *sativus*, we compared the transcribed sORFs found in different tissues using a Venn diagram (Figure 2). For Jin You 1 cultivar, a total of 1,442 (27.74%) transcribed sORFs were expressed in both root and leaf tissues, and 2,681 and 1,076 transcribed sORFs were only expressed in the root or leaf tissue, respectively (Figure 2a). For Jin You 35 cultivar, 5,131 transcribed sORFs were expressed in both leaf and root tissues, and 2,322 and 3,661 transcribed sORFs were specifically expressed in root and leaf tissues, respectively (Figure 2b). Results from the Venn diagram analysis showed only a total of 83 (0.59%) transcribed sORFs were shared among the different tissues of cv. Chinese Long (Figure c). Tissue-specific transcribed sORFs were also detected in Chinese Long cultivar: vascular tissues (534), fruit flesh (701), root (236), shoot apex (169), leaf (3,247). As for the Zaoer-N cultivars (Figure 2d), a total of 2,454 transcribed sORFs were detected in all three tissues: epidermis, basic tissue, and vascular bundle. Tissue-specific transcribed sORFs were also detected in Zaoer-N cultivar: epidermis (1,404), basic tissue (988), and vascular bundle (1,725).

**Figure 1:**
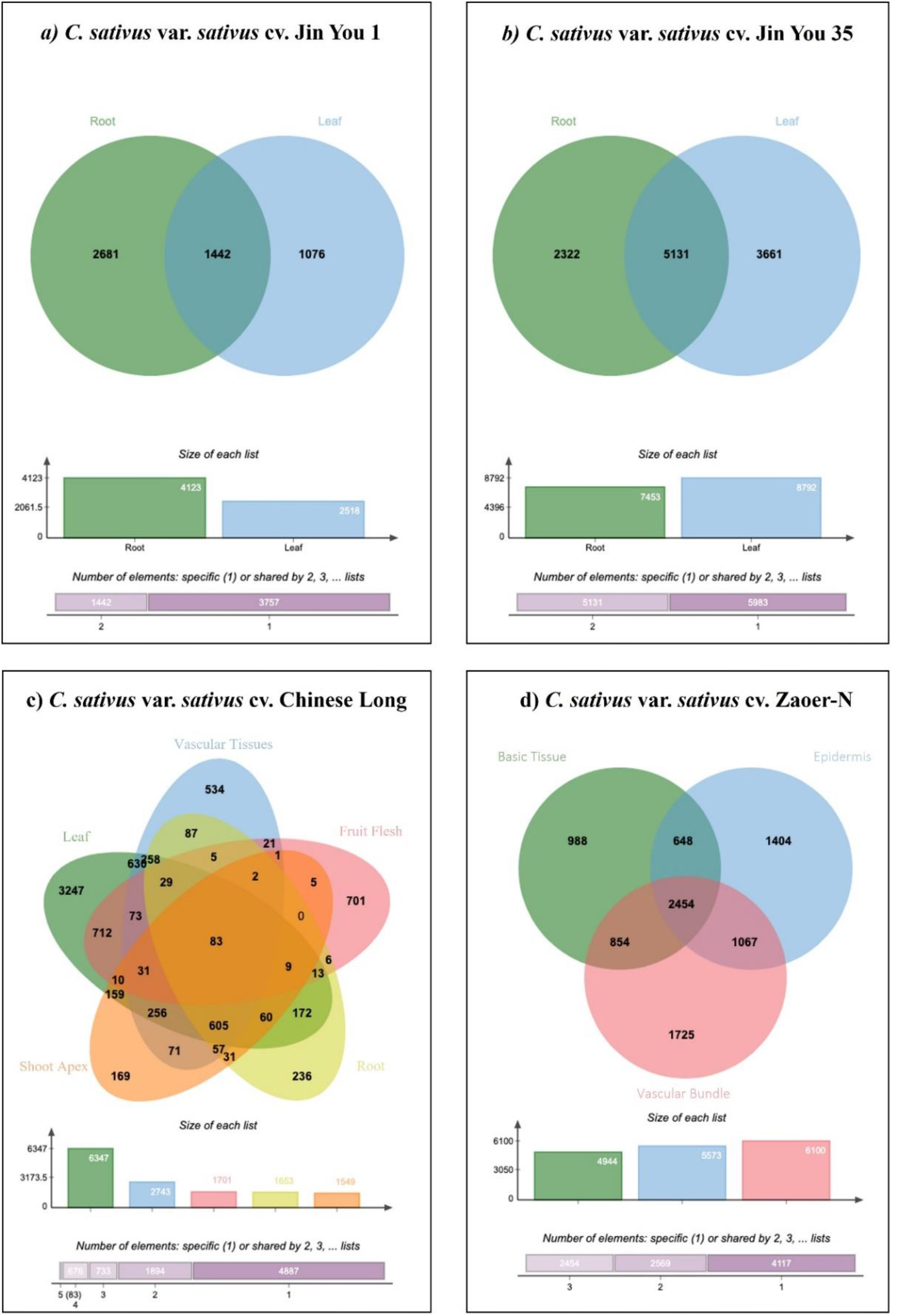
Comparison of the transcribed sORF in transcriptome datasets of different tissue samples: a) Jin You 1 cultivar, b) Jin You 35, c) Chinese Long and d)Zaoer-N.

**Figure 2:**
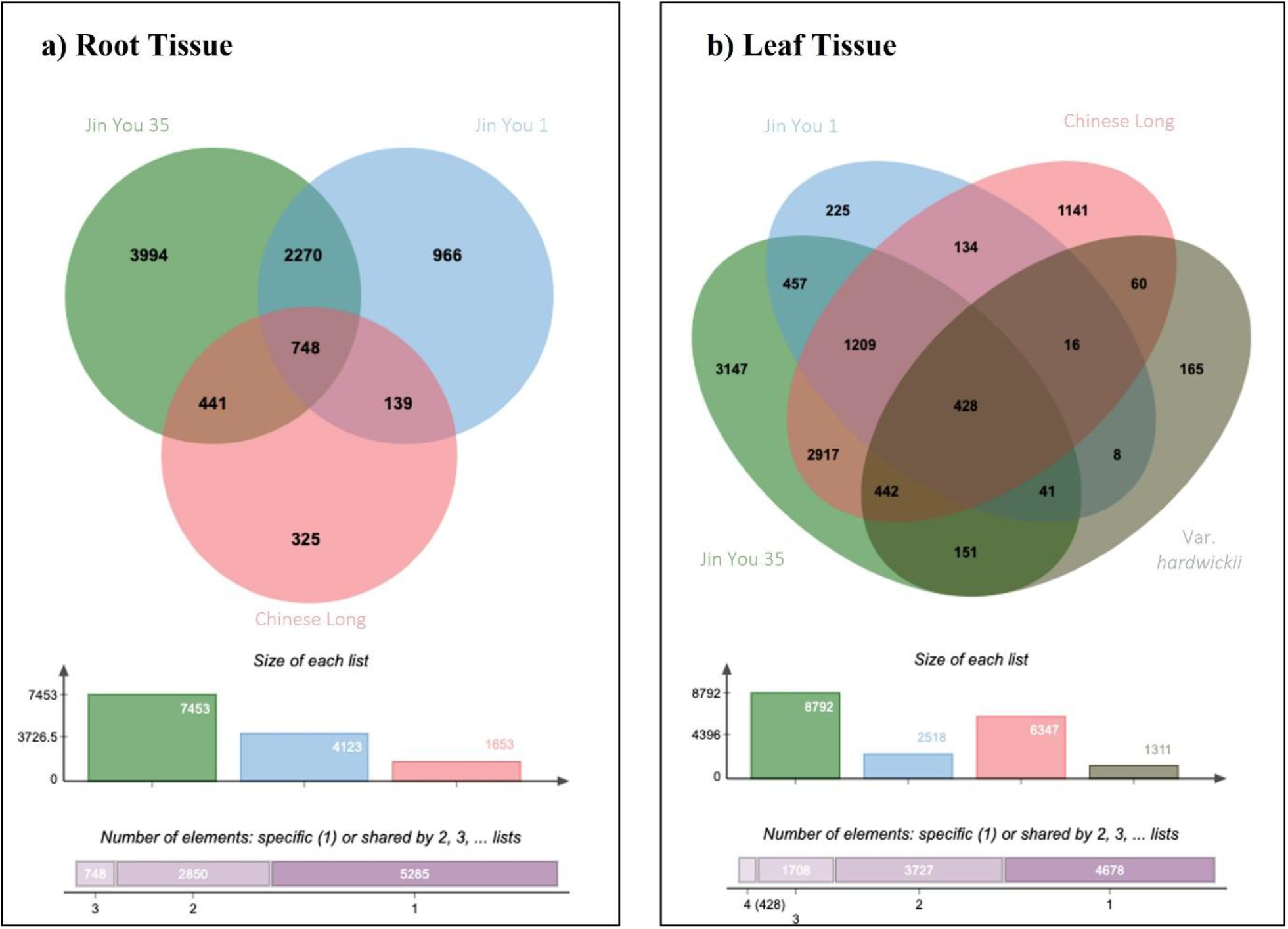
Comparison of the transcribed sORF in root (a) and leaf (b) transcriptome datasets of different *C. sativus* var. *sativus* cultivars.

The transcribed sORFs in the root tissue of three cultivars of *C. sativus* var. *sativus* were compared in this study (Figure 3a). Only 748 (8.42%) transcribed sORFs were shared among the root tissues of different cultivars. Overall, the root tissue of cv. Jin You 35 has more transcribed sORFs (3,994) compared to cv. Jin You 1 (966) and cv. Chinese Long (325). A total of 748 and 428 high-confident root tissue-specific and leaf tissue-specific transcribed sORFs were identified in different cultivars respectively (Figure 3). For transcribed sORF in leaf tissues of 4 cultivars, cv. Jin You 35 topped the chart with almost 9,000 transcribed sORFs detected, followed by Chinese Long 9930 (5,697), Jin You 1 (2,518), and lastly var. *hardwickii* (1,311). In *A. thaliana*, Hanada et al. (2013) generated a coding sORF expression atlas for 16 organs and 17 environmental conditions and showed that 2,099 transcribed sORFs were highly expressed under at least one environmental condition and 571 transcribed sORFs were significantly conserved in other plant species.

**Figure 3:**
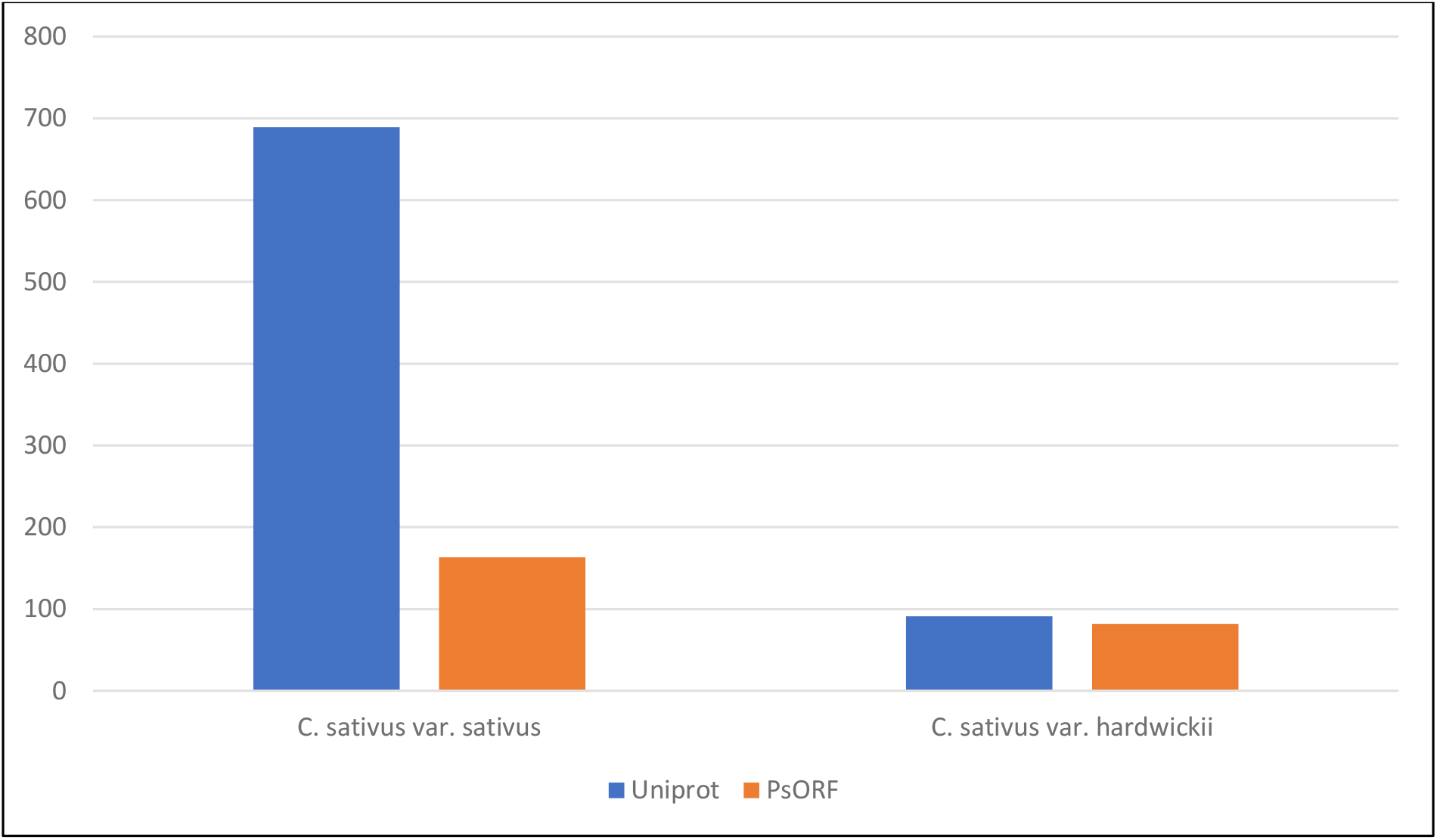
Transcribed sORFs with translational potential in *C. sativus* var. *hardwickki* and var. *sativus*.

### Transcribed sORF with translational potential in *Cucumis sativus*

Using a protein sequence homology search approach, only 689 (4.63%) out of the 14,866 transcribed sORFs found in *C. sativus* var. *sativus* might have translational potential (Figure 4). *C. sativus* var. *hardwickii* on the other hand has a slightly higher percentage of transcribed sORFs with translational potential (6.94%; 91 translated sORFs) compared to var. *sativus*. Besides SWISS-PROT, we also blast searched the transcribed sORFs against the sORFs with translational potential deposited in the PsORF database. The PsORF database is a collection of plant sORFs from 35 different plant species (Chen et al., 2020). The authors collected multi-omics datasets including genome, transcriptome, Ribo-seq, and mass spectrum from public databases and built a bioinformatics pipeline to detect sORFs in these datasets. Results from blast search against the PsORF database showed that 91.23% (*C. sativus* var. *sativus*) and 90.11% (*C. sativus* var. *sativus*) of transcribed sORFs with the homologs in SWISS-PROT also have homologs in the PsORF database (Figure 4). This indicates that these transcribed sORFs might have translational potential. Using a proteomic approach, Castellana et al. (2008) identified ~5,000 small peptides in Arabidopsis and some of these small peptides were novel and/or identified by Hanada et al. (2007).

**Figure 4:**
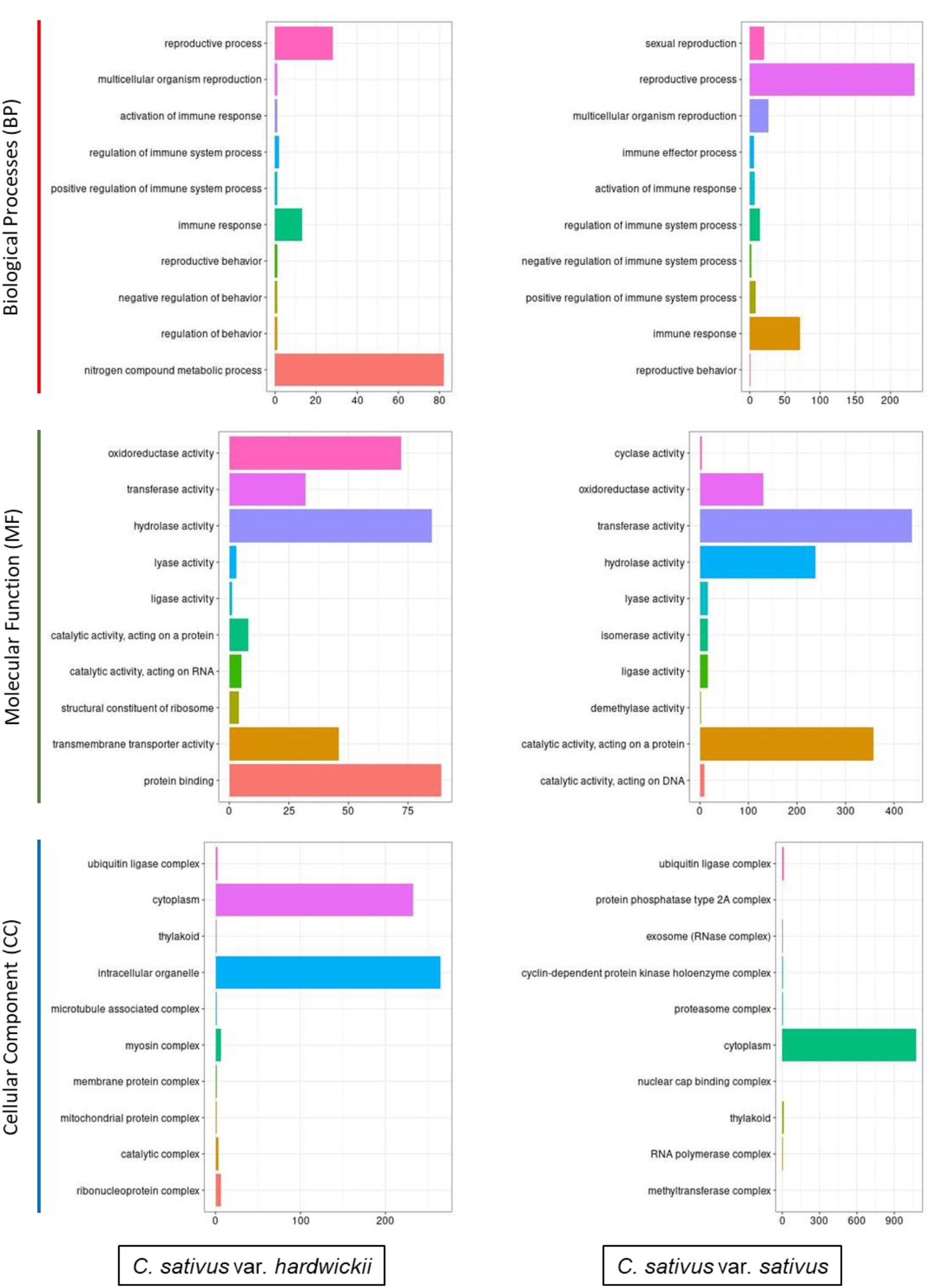
Gene ontology (GO) classification of *C. sativus* var. *hardwickii* and *C. sativus* var. *sativus* transcribed sORFs.

### Functional Classification of *Cucumis sativus* sORF

From the 689 and 91 transcribed sORFs with the translational potential of *C. sativus* var. *hardwickii* and var. *sativus* respectively, 2,005 and 6,884 unigenes with Entrez ID were retrieved from the UnitProt database for Gene Ontology (GO) analysis using the clusterProfiler (Figure 5). Among the 3 distinct categories of GO classes, molecular functions were the most represented functional group in both varieties for sORF functional annotation (Figure 5). The transcribed sORFs involved in BP category of C. *sativus* var. *hardwickii* and var. *sativus* share 7 GO terms. These GO terms are reproductive process, multicellular organism reproduction, activation of immune response, regulation of immune system process, regulation of immune system process, immune response, and reproductive behaviour. Overall, the number of the transcribed sORFs in the 7 shared GO terms is higher in C. *sativus* var. *sativus* as compared to var. *hardwickii* except for the reproductive behaviour term (Figure 5). As for the MF and CC categories, both varieties shared 6 and 3 GO terms, respectively.

**Figure 5:**
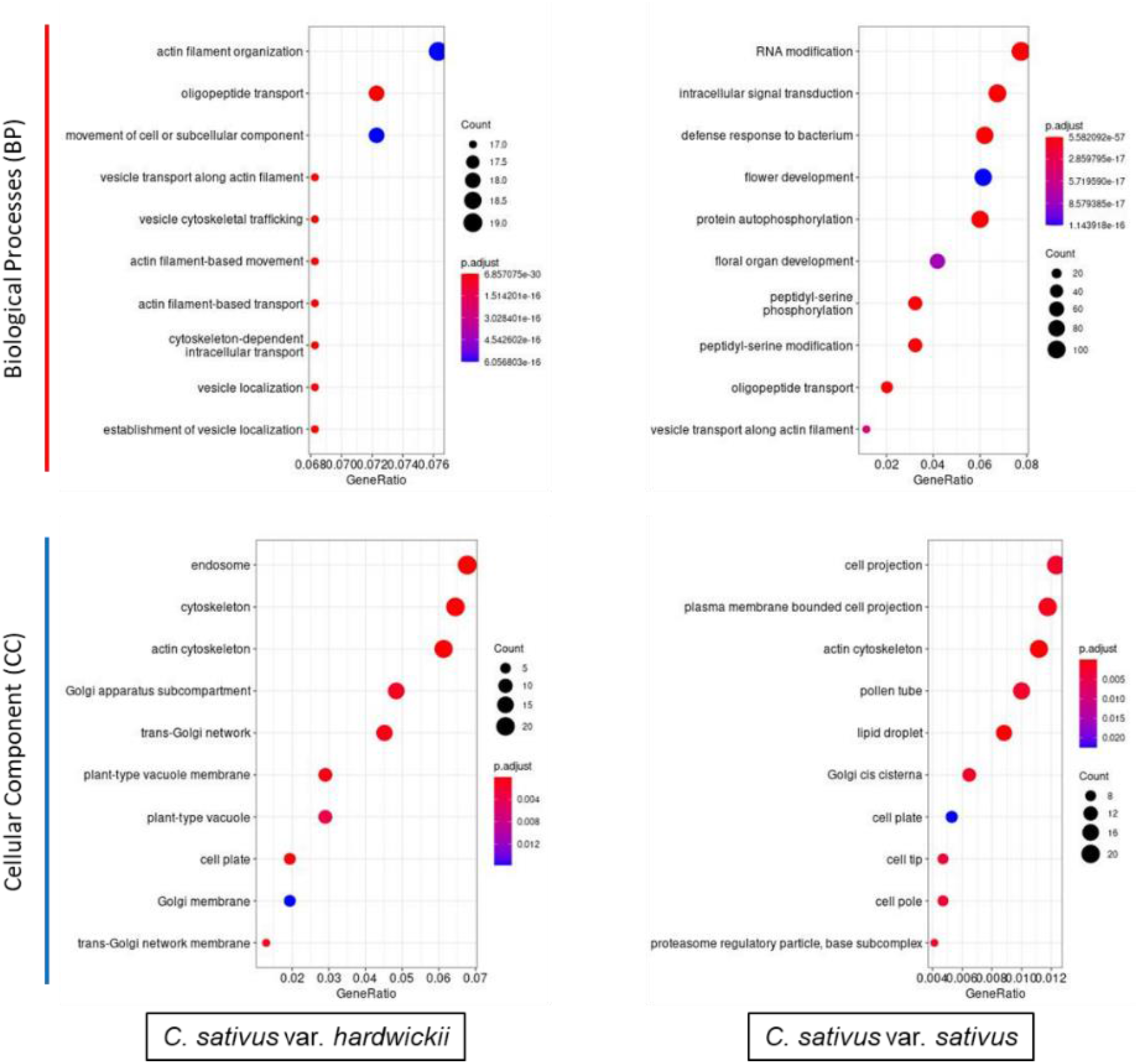
Bubble plot showing the enriched GO terms. X-axis in the bar plot stood for gene ratio, while the y-axis indicates different BP and CC. The size of the circles in each plot is positively correlated with the number of genes involved in each subgroups while the colour of the circles indicate their significance level.

We also performed the GO enrichment analysis of transcribed sORFs of both varieties (Figure 6). The transcribed sORFs were enriched in the GO terms of BP and CC. No enrichment was detected for MF. In C. *sativus* var. *hardwickii*, most of the transcribed sORFs showed significant CC enrichment in the endosome, cytoskeleton, and actin cytoskeleton and to a lesser extent in the Golgi membrane and trans-Golgi network membrane. The cytoskeleton mainly functions as a structure for cell shape and internal organization and is made up of 3 elements, namely, microtubules, intermediate filaments, and actin (Microtubules and Filaments, 2014). As shown in the GO enrichment plot (Figure 6), the transcribed sORFs with enriched GO terms related to actin were detected in both varieties. Actin mainly functions in cellular physiological processes in plants including cell growth, cytokinesis, cell division, and several intracellular trafficking events (Diao and Huang, 2021). This indicates that the transcribed sORFs of *C. sativus* might play a role in growth and development regulations. Hanada et al. (2013) reported that overexpression of sORFs showed varying morphological changes in transgenic *A. thaliana* and was associated with a higher growth rate.

For the enriched KEGG pathways, the transcribed sORFs in both varieties were enriched in KEGG Ontology (KO) terms of plant hormone signal transduction. Plant sORFs have been demonstrated to play important roles in cell signalling, abiotic stress response, morphogenesis, and growth regulation (Hanada et al., 2013; Bashir et al., 2014; Ong et al., 2022). Apart from that, some of the transcribed sORFs identified in *C. sativus* var. *sativus* are significantly associated with the plant-pathogen interaction. This indicates that *C. sativus* transcribed sORFs play a significant role in plant growth and development, and environmental stress responses.

## 3. Material and Methods

### 3.1 Data retrieval of Cucumis sativus reference genome and transcriptomes

The reference genome sequences and annotation file of *C. sativus* var. *hardwickii* PI183967 were retrieved from CuGenDBv2 (http://cucurbitgenomics.org/). The transcriptome datasets of *C. sativus* var. *hardwickii* and *C. sativus* var. *sativus* were retrieved from NCBI SRA database (https://www.ncbi.nlm.nih.gov/sra). The details of the retrieved transcriptome datasets were summarised in Supplementary Table 1.

### 3.2 In silico prediction of small open reading frame

The CDS, exon, intron, and intergenic regions of *C. sativus var. hardwickii* PI183967 were extracted from its reference genome using gff2sequence (Camiolo and Porceddu, 2013). sORFfinder (Hanada et al., 2010) was used to predict the coding sORFs from *C. sativus var. hardwickii* PI183967 genome. RepeatMasker (http://www.repeatmasker.org) was used to mask the repeat sequences in the coding sORFs and the masked sequences were removed using an in-house script. Sequence clustering of the coding sORFs was performed using CD-HIT (Li and Godzik, 2006) with a clustering threshold of 95% to cluster the redundant sequences into sequence clusters.

### 3.3 Characterisation of small open reading frame

To identify transcribed sORFs, the coding sORFs were blast searched against Cucumber EST collection version 3 (http://cucurbitgenomics.org/est/cucumber) using the blastn algorithm with homology search parameters “-evalue 1e-5 -per_identity 97 - qcov_hsp_perc 100”. The nucleotide sequences of transcribed sORFs were translated to amino acid sequences using gotranseq (https://github.com/feliixx/gotranseq). The amino acid sequences of transcribed sORFs were blast searched locally against the high quality manually annotated and non-redundant protein sequences retrieved from the SWISS-PROT database (https://www.uniprot.org/help/downloads) using the blastp algorithm with homology search parameters “-evalue 1e-5 -per_identity 97 - qcov_hsp_perc 100”. The transcribed sORFs that shared high homology to SWISS-PROT protein sequences were designated as transcribed sORFs with translation potential (translated sORFs). The amino acid sequences of translated sORFs were then blast searched against the plant sORF database, PsORF (http://psorf.whu.edu.cn/; Chen et al., 2020).

### 3.4 Transcriptome analysis of small open reading frames

The csORF sequences of *C. sativus* var. *hardwickii* PI183967 were mapped to its reference genome using blat (https://github.com/djhshih/blat) and the outputs were converted to gene annotation file using blat2gtf.pl (https://github.com/IGBIllinois/HOMER/blob/master/bin/blat2gtf.pl). The sORF annotation file was combined with the reference genome annotation file using agat_sp_merge_annotations.pl from AGAT package (https://github.com/NBISweden/AGAT). The RNA-seq reads were aligned and mapped to *C. sativus var. hardwickii* PI183967 genome using HISAT2 (http://daehwankimlab.github.io/hisat2/) and the expression profile of sORFs in the RNA-seq datasets were determined using Stringtie (https://ccb.jhu.edu/software/stringtie/#install) together with the updated gene annotation file. The sORF sequence IDs were retrieved from the Stringtie output file using an in-house script and the sORF sequences were then extracted from the coding sORF file using the subseq function in the seqtk package (https://github.com/lh3/seqtk). The sORF sequences identified by HISAT2/Stringtie pipeline were combined with the sORF sequences that shared homology to cucumber ESTs to form a final set of transcribed sORF. The transcribed sORF sequences that can be found in both *C. sativus* var. *hardwickii* and *C. sativus* var. *sativus* RNA-seq datasets were identified using online Venn diagram plotters (https://bioinformatics.psb.ugent.be/webtools/Venn/; http://jvenn.toulouse.inra.fr/app/example.html, Bardou et al., 2014) and designated as high confident species-specific transcribed sORF. The unique transcribed sORFs from each variety were designated as varietal-specific transcribed sORF.

### 3.5 Functional annotation of small open reading frame

To assign a biological function to transcribed sORF, Gene Ontology (GO) and the Kyoto Encyclopedia of Genes and Genomes (KEGG) were performed using the clusterProfiler R package (https://bioconductor.org/packages/release/bioc/html/clusterProfiler.html). The transcribed sORF amino acid sequences were first blast searched against SWISS-PROT database using the blastp algorithm. The SWISS-PROT protein IDs were extracted from the blast output using an in-house script and converted to GeneID using the “Retrieve/ID Mapping” function from the UniProt website (https://www.uniprot.org/id-mapping). The GO category analysis was conducted using the groupGO function embedded in the clusterProfiler R package. The enrichment analyses of GO and KEGG were performed using the enrichGO and enrichKEGG function embedded in the clusterProfiler R package.

## 4. Conclusion

In this study, we have established a bioinformatics pipeline for the identification of small open reading frames (sORFs) in *Cucumis sativus*. Using the pipeline, different types of sORFs were identified from the genome and transcriptome datasets of *Cucumis sativus* var. *hardwickii* and var. *sativus*. GO and KEGG terms that enriched in growth and development, and stress response were predicted for the transcribed sORFs with translational potential. Further classification of sORFs is needed to minimise conflicting sORF annotations and ease categorisations of sORFs. Having a complete database of transcribed sORFs and translated sORFs in *Cucumis sativus* would help us understand the roles of plant SEPs, especially in the biotic and abiotic stress responses. With that being said, cucumber producers will be able to improve crop viability, especially in harsh weather conditions, and produce cucumbers with higher market value.

## Author Contributions

Conceptualization, B.C.T, C.H.T; experimental design, G.S.W.C, C.H.T; bioinformatics analysis, G.S.W.C, C.H.T; manuscript writing, G.S.W.C, B.C.T, C.H.T. All authors have read and agreed to the published version of the manuscript.

