## Supplementary Table 1 for "*In silico* comparative RNA-seq analysis reveals varietal-specific intergenic small open reading frames in *Cucumis sativus* L."

**Supplementary Materials**

**Supplementary Table 1:** RNA-seq datasets used.

| No. | Cultivar | Tissue | Accession number | Number of reads |
|---|---|---|---|---|
| 1 | Jin You 1 | Root | SRR8372183<br>SRR8372185<br>SRR8372186 | 35605649 |
| | | Leaf | SRR7429308<br>SRR7429309<br>SRR7429310 | 35605638 |
| 2 | Jin You 35 | Root (12hr) | SRR8662481<br>SRR8662478<br>SRR8662479 | 57496572 |
| | | Root (24hr) | SRR8662490<br>SRR8662489<br>SRR8662485 | 52043451 |
| | | Leaf (12hr) | SRR8662510<br>SRR8662509<br>SRR8662512 | 60560232 |
| | | Leaf (24hr) | SRR8662497<br>SRR8662496<br>SRR8662495 | 77975756 |
| | | Leaf | SRR18498271<br>SRR18498272<br>SRR18498273<br>SRR18498274<br>SRR18498275 | 111899794 |
| 3 | Chinese Long | Internode vascular tissue | SRR3156393<br>SRR3156394<br>SRR3156395 | 15147778 |
| | | Lamina | SRR3156399<br>SRR3156400<br>SRR3156401 | 15147778 |
| | | Major Vein | SRR3156405<br>SRR3156406<br>SRR3156407 | 38362588 |
| | | Phloem Sap | SRR3156411<br>SRR3156412<br>SRR3156413 | 35756932 |
| | | Petiole Vascular Tissue | SRR3156417<br>SRR3156418<br>SRR3156419 | 39832709 |
| | | Root | SRR3156423<br>SRR3156424<br>SRR3156425 | 34981075 |
| | | Shoot Apex | SRR3156429<br>SRR3156430<br>SRR3156431 | 32754627 |

| | | | | |
|---|---|---|---|---|
| | | Fruit Flesh | SRR5405124<br>SRR5405125 | 11249494 |
| | | Leaf | SRR6895190<br>SRR6895191<br>SRR6895192 | 98752091 |
| 4 | Zaoer-N | Epidermis | SRR19513186<br>SRR19513201<br>SRR19513200 | 67915646 |
| | | Basic Tissue | SRR19513196<br>SRR19513195<br>SRR19513194 | 57437754 |
| | | Vascular Bundle | SRR19513197<br>SRR19513198<br>SRR19513199 | 69413495 |
| 5 | Hardwickii | Leaf | SRR11573113 | 36560996 |
